# Mouse Norovirus infection arrests host cell translation uncoupled from the stress granule-PKR-eIF2α axis

**DOI:** 10.1101/536052

**Authors:** Svenja Fritzlar, Turgut E. Aktepe, Yi-Wei Chao, Michael R. McAllaster, Craig B. Wilen, Peter A. White, Jason M. Mackenzie

**Author notes:** Present address: Department of Microbiology, Biomedical Discovery Unit, Monash University, Melbourne, Victoria, Australia (Svenja Fritzlar). Corresponding author: (JMM). These authors contributed equally to this work. SF and TEA are Joint Authors.

## Abstract

The integrated stress response (ISR) is a cellular response system activated upon different types of stresses, including viral infection, to restore cellular homeostasis. However, many viruses manipulate this response for their own advantage. In this study we investigated the association between murine norovirus (MNV) infection and the ISR and demonstrate that MNV regulates the ISR by activating and recruiting key ISR host factors. We observed that during MNV infection, there is a progressive increase in phosphorylated eukaryotic initiation factor 2 alpha (p-eIF2α) resulting in the suppression of host translation, yet MNV translation still progresses under these conditions. Interestingly, the shutoff of host translation also impacts the translation of key signalling cytokines such as IFNβ, IL-6 and TNFα. Our subsequent analyses revealed that the phosphorylation of eIF2α was mediated via Protein kinase-R (PKR), but further investigation revealed that PKR activation, phosphorylation of eIF2α and translational arrest were uncoupled during infection. We further observed that stress granules (SGs) are not induced during MNV infection, and MNV has the capacity to restrict SG nucleation and formation. We observed that MNV recruited the key SG nucleating protein G3BP1 to its replication sites and intriguingly the silencing of G3BP1 negatively impacts MNV replication. Thus, it appears, MNV utilises G3BP1 to enhance replication, but equally to prevent SG formation, intimating an anti-MNV property of SGs. Overall, thus study highlights MNV manipulation of SGs, PKR and translational control to regulate cytokine translation and to promote viral replication.

**Importance:** Viruses hijack host machinery and regulate cellular homeostasis to actively replicate their genome, propagate and cause disease. In retaliation, cells possess various defence mechanisms to detect, destroy and clear infecting viruses as well as signal to neighbouring cells to inform them of the imminent threat. In this study, we demonstrate that the murine norovirus (MNV) infection stalls host protein translation and the production of antiviral and pro-inflammatory cytokines. However, virus replication and protein translation still ensues. We show that MNV further prevents the formation of cytoplasmic RNA granules, called stress granules (SG), by recruiting the key host protein G3BP1 to the MNV replication complex; a recruitment that is crucial to establishing and maintaining virus replication. Thus MNV promotes immune evasion of the virus by altering protein translation. Together, this evasion strategy delays innate immune responses to MNV infection and accelerates disease onset.

## Introduction

Human noroviruses (HuNoV) are positive sense single-stranded RNA viruses and belong to the *Caliciviridae* family. They are a major cause of acute gastroenteritis in developing and developed countries (1–3). The onset of symptoms like diarrhoea, nausea, vomiting and abdominal cramps usually commences 12-48 hours after exposure to the virus and typically lasts no more than 48 hours (4–6). Despite its significant health burden, there are currently no effective treatments or preventative vaccines for HuNoV infections even though vaccines are under development (7–11). Advances of antiviral agents to control HuNoV outbreaks are severely delayed by the fact that HuNoVs are difficult to cultivate in the laboratory. Recent studies have shown that HuNoV is able to replicate in B-cell like cell lines when co-cultured with specific enteric bacteria or in enteric organoids (12, 13). However viral replication is poor with only a 2-3 Log increase in viral titre and therefore, the closely related Genogroup V murine norovirus (MNV) remains a robust tissue culture system and small animal model (14).

The MNV genome is a ~7.5 kb positive-sense RNA molecule that encodes for 9 or 10 proteins (depending on translation of open reading frames (ORFs) and cleavage of gene products (15, 16)); that have roles in replication of the viral genome, polyprotein cleavage, translation, host manipulation and assembly of virus particles. The genome itself is covalently attached to the viral protein g (VPg or NS5) at its 5’ end and is polyadenylated at the 3’ end. The VPg protein mediates translation of the viral genome via interaction with host translation factors (17, 18). The remaining non-structural proteins (ORF1) associate with the viral replication complex (RC) in induced membrane clusters (19, 20), as well as interacting with host factors to manipulate cellular homeostasis and promote viral replication. Not all proteins encoded by ORF1 have been functionally characterised, but previous studies revealed that the MNV NS1/2 protein associates with the ER and the host protein VAP-A (21, 22), whereas NS3 associates with microtubules and lipid rich bodies in the cytoplasm (23). Further, NS7 acts as the RdRp (24, 25) and NS6 is the protease cleaving the polyprotein (26, 27).

Noroviruses cause acute and chronic infections that often involve manipulation of host processes and innate immune responses at multiple levels (Reviewed in 28). The introduction of viral dsRNA and proteins during MNV infection is recognised as foreign by the integrated stress response (ISR), and this can activate antiviral innate immune pathways. This recognition of infection can result in a myriad of responses with the most important being the type I and type III interferon (IFN) response (29), however the exact mechanisms employed to restrict and clear a NoV infection are not completely defined

In the presence of cellular stressors, the ISR can be activated by eIF2α kinases such as double-stranded RNA sensor PKR, the ER-stress sensor PKR-like endoplasmic reticulum kinase (PERK), general control non-derepressible 2 kinase (GCN2) and heme-regulated kinase (HRI). The activation of these sensors can lead to the phosphorylation of eIF2α which relinquishes eIF2α’s ability to bind to the 40s ribosomal subunit, prompting translational stalling and the aggregation of stalled translation preinitiation complexes (30–32). Together with Ras-GAP SH3 domain binding protein (G3BP), T-cell restricted intracellular antigen 1 (TIA-1) and TIA-1 related protein (TIAR), these aggregates form stress granules (SGs) to stall translation and protect the cell from accumulating misfolded proteins during viral infections. SGs contain the preinitiation complexes which are typically comprised of various initiation factors including eIF2, eIF3, eIF4α, eIF4β, eIF4G and eIF5, as well as the 40s ribosomal subunit which ensures that SGs reactivate translation rapidly after a successful stress recovery (33). SGs have also been shown to regulate and control cytokine mRNA aggregation and expression.

Several viruses manipulate the ISR to avoid immune detection by inhibiting translation and preventing the formation of SGs. Sinbis virus strongly inhibits the translation of cellular mRNA in PKR-dependent, as well as PKR-independent mechanisms (34). Influenza A virus inhibits the phosphorylation of eIF2α and therefore prevents the induction of stress granules (35). Poliovirus, Herpes simplex virus 1 and West Nile virus (WNV) interfere with SG formation by cleaving or sequestering SG nucleating proteins like G3BP and TIA-1 (36–38).

In this study we demonstrate that MNV infection leads to the phosphorylation of eIF2α, via PKR, and the subsequent host cell translational shutoff but this does not affect viral translation. However, restoration of active eIF2α does not alleviate host cell translational repression suggesting that these events are uncoupled. Further, we show that this translational shut-off is associated with a decrease in cytokine translation and that MNV inhibits the formation of SGs by recruiting the SG nucleating factor G3BP1 to the sites of virus replication, a recruitment that is essential for MNV replication. Thus, we provide evidence that MNV manipulates the PKR–p-eIF2α–SG axis to promote its own replication but equally as an immune evasion strategy.

## Results

### MNV infection induces eIF2α phosphorylation

During our investigations of the intracellular replication of MNV we noticed changes in the abundance of host cell protein translation. To initially interrogate the influence of MNV on translation, we investigated whether MNV infection and replication induced eIF2α phosphorylation (Fig. 1 A and B). BMM cells were left untreated (mock), treated with the oxidative stressor sodium arsenite (NaAs; 250 μM for 20 minutes (mins)) or infected with MNV for 12 hours (hrs). Our western blot (WB) analysis of whole cell lysates revealed that MNV infection, similar to the NaAs positive control, induced a robust increase in p-eIF2α levels, whereas the total levels of eIF2α remained constant (Fig. 1A).

**Figure 1.**
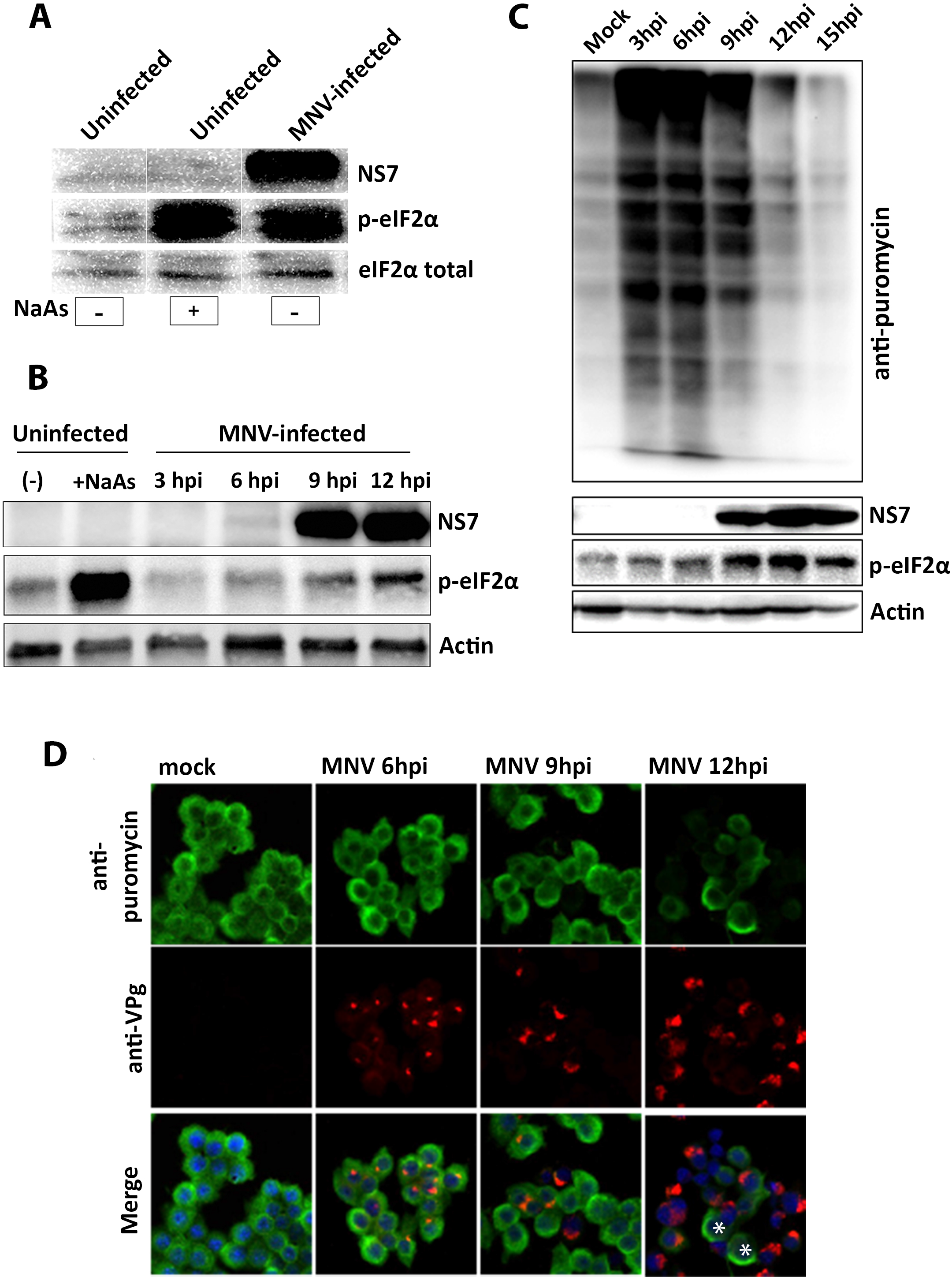
MNV infection phosphorylates eIF2α and shuts down host cell translation: **(A)** BMMs were uninfected, uninfected and NaAs treated (250 μM for 20 mins) or MNV-infected (MOI 5) for 12 hrs. The WB was immunolabelled with anti-NS7, anti-p-eIF2α and anti-eIF2α antibodies **(B)** Immunoblot analysis of uninfected, uninfected and NaAs treated (250 μM for 20 mins) or MNV-infected (MOI 5) cell lysates harvested at 3, 6, 9 and 12 h.p.i. The WB was immunolabelled with anti-NS7, anti-p-eIF2α and anti-actin antibodies. **(C and D)** BMM cells were either infected with MNV (MOI 5) or left uninfected and analysed for their translation using puromycin (10 μg/mL). (**C**) Immunoblot analysis of puromycin-treated (20 mins) cell lysates harvested at 3, 6, 9, 12 and 15 h.p.i. The WB was immunolabelled with anti-puromycin, anti-NS7, anti-p-eIF2α and anti-actin antibodies. (**D**) IF analysis of puromycin treated (10 μg/ml for 30 mins) cells at 6, 9 and 12 hrs post infection. Cells were stained with anti-puromycin, anti-NS5 and DAPI for the merged image. Stars indicate uninfected cells displaying high signal for anti-puromycin. Samples were analysed via the Zeiss LSM 710 confocal microscope and analysed with the ZEN software.

To further these initial observations, we investigated the kinetics of eIF2α phosphorylation throughout the course of the viral infection. Thus, MNV-infected cell lysates were collected at 3, 6, 9 and 12 hours post infection (hpi) and WB analysis was performed with antibodies for p-eIF2α, MNV NS7 and actin (Fig. 1B). Interestingly, we observed that eIF2α phosphorylation levels remained constant at 3 hpi when compared to uninfected and untreated cells, however as infection progressed, there was a gradual and noticeable increase in eIF2α phosphorylation at 6, 9 and 12 hpi. This indicates that MNV infection induces increased phosphorylation of eIF2α as infection proceeds (Fig. 1B).

### eIF2α phosphorylation status corresponds to a repression of host cell protein translation during MNV infection

One of the main consequences of eIF2α phosphorylation is the global shutoff of host cell protein translation (39, 40). To determine the effects of increasing eIF2α phosphorylation levels on cellular translation, BMM cells were infected with MNV, and at indicated times post-infection (3, 6, 9, 12 and 15 hpi), cells were pulsed with puromycin for 20 mins prior to whole cell lysate collection (for WB) and cell fixation (for immunofluorescence (IF)) (Fig. 1C and D, respectively).

Puromycin incorporates into newly translated polypeptides and terminates the translation of the full-length protein. Thus, newly synthesised proteins, which accurately represents translational activity, can therefore be visualised using an anti-puromycin antibody. Our WB analysis revealed that there was an increase in the amount of puromycin incorporated in active protein translation up to 6 hpi (Fig. 1C). However, as the infection progressed from 9 hpi (indicative by NS7 labelling) and the level of p-eIF2α increased, the levels of puromycin-labelled proteins were reduced (Fig 1C). These results were supported by IF analysis demonstrating that incorporation of puromycin (Fig 1D, green) was significantly diminished in MNV-infected cells from 9 hpi (Fig. 1D), whereas the uninfected mock cells still incorporated substantial amounts of puromycin (Fig. 1D). These results confirm that the MNV-induced increase in eIF2α phosphorylation results in decreased host cell protein translation. Interestingly, MNV protein translation (as determined by NS7 expression) steadily increases over the course of the infection even in the presence of eIF2α phosphorylation and host cell protein translation shut down (Fig. 1C).

### PKR induces the phosphorylation of eIF2α during MNV infection, but translation repression is PKR-independent

During viral infection, PKR and PERK are two major kinases induced to prevent viral replication by phosphorylating eIF2α and inhibiting translation (41, 42). To investigate the potential role of PKR and/or PERK in mediating phosphorylation of eIF2α during MNV infection, we utilised a PKR inhibitor (C16, shown to suppress PKR-mediated phosphorylation of eIF2α (43) and a PERK inhibitor (ISRIB, shown to suppress PERK-mediated phosphorylation of eIF2α (44) to treat MNV-infected RAW 264.7 cells (Fig. 2A). Cells were infected with MNV and subsequently treated with 1 μM C16 and/or 0.5 μM ISRIB at 1 hpi. Cell lysates were collected at 12 hpi, and WB analysis was performed using an anti-p-eIF2α antibody. C16 treatment of MNV infected cells substantially decreased the levels of p-eIF2α compared to untreated MNV infected cells. In contrast, we observed no apparent change in the p-eIF2α levels in the ISRIB treated MNV-infected cells. These results indicate that the phosphorylation of eIF2α observed during MNV infection is primarily mediated via PKR and not PERK.

**Figure 2.**
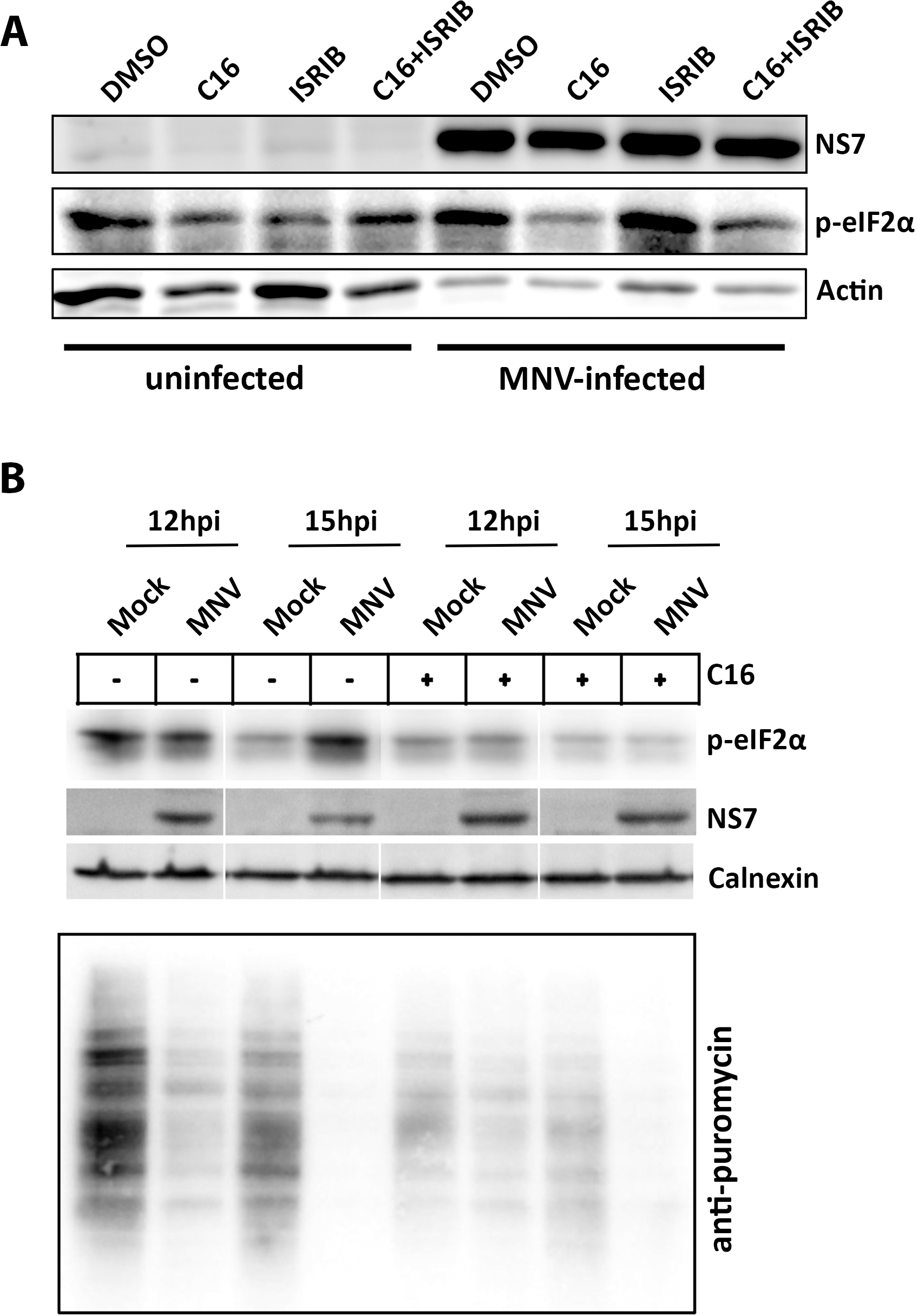
Treatment with PKR inhibitor C16 abolishes phosphorylation of eIF2α but does not rescue host translation: **(A)** RAW 264.7 were either uninfected or MNV-infected (MOI 5), treated with either DMSO, C16 (1 μM), ISRIB (0.5 μM) or C16+ISRIB at 1 h.p.i for 12 hrs, before cell lysate samples were obtained. Lysates were analysed via immunoblotting and immunolabelled with anti-NS7, anti-p-eIF2α or anti-actin antibodies. **(B)** RAW 264.7 were either uninfected or MNV-infected (MOI 5), treated with either DMSO or C16 (1 μM) at 1 h.p.i for 12 or 15 hrs. 30 mins before cell lysate samples were obtained, cells were treated with puromycin (10 μg/ml for 30 mins) and immunolabelled with anti-NS7, anti-p-eIF2α, anti-puromycin and anti-calnexin antibodies.

Due to our observations that eIF2α phosphorylation was mediated via PKR, we speculated that inhibition of PKR activity would restore the repression of host cell protein translation. Following MNV infection and C16 treatment, cells were incubated with puromycin before harvesting lysates at 12 or 15 hpi. Similar to previous results, protein translation is severely inhibited in the untreated MNV-infected cells, however surprisingly this phenotype was also maintained even in the presence of C16 and the lack of p-eIF2α (Fig. 2B). Thus, our results indicate that MNV-induced repression of host translation is uncoupled and independent of a PKR and p-eIF2α-mediated mechanism and must occur via a different regulatory pathway.

### The MNV NS3 protein induces host cell protein translation arrest

Previous studies have suggested that the MNV protease NS6 can influence host cell protein translation via cleavage of the translation accessory factor PABP (45). To verify these observations, we expressed PABP-GFP in RAW 264.7 and infected the cells with MNV for 12 h. Neither the viral protein NS5 (VPg) nor NS6 (Protease) co-localised with PABP-GFP in infected cells and PABP-GFP expression was still observed (Fig. S1A). Further, we co-transfected cDNA expression plasmids encoding MNV NS3, NS6 or NS7 (RdRp) with PABP-GFP in HEK 293T cells. Upon co-transfection with NS6 and NS7, PABP-GFP expression levels seemed unperturbed, however co-transfecting PABP-GFP with NS3 significantly reduced PABP-GFP levels compared to the control (Fig. S1B). It is important to note that we did not detect any smaller sized protein bands for PABP-GFP that would indicate virus-induced cleavage of this protein, even in the presence of the MNV NS6 protease expressed during replication (Fig. S1B).

To interrogate which MNV proteins might affect host cell translation, we utilised puromycin incorporation in individual ORF1 protein transfected cells. Thus, HEK 293T and Vero cells were transfected with plasmids encoding the single ORF1 proteins and treated with puromycin prior to immunoblot or IF analysis. We observed no significant change in the amount of incorporated puromycin in cells expressing MNV NS1/2, NS4, NS5, NS6 or NS7. Intriguingly, we observed a profound absence of puromycin incorporation in cells expressing the MNV NS3 protein (Figs. 3A, B and C). These results suggest that the MNV NS3 has an impact on the host protein translational efficiency. In contrast to previous reports, we observed that the MNV NS6 protease did not influence host protein translation, nor cleave the translation accessory factor PABP (Fig. S1).

**Figure 3.**
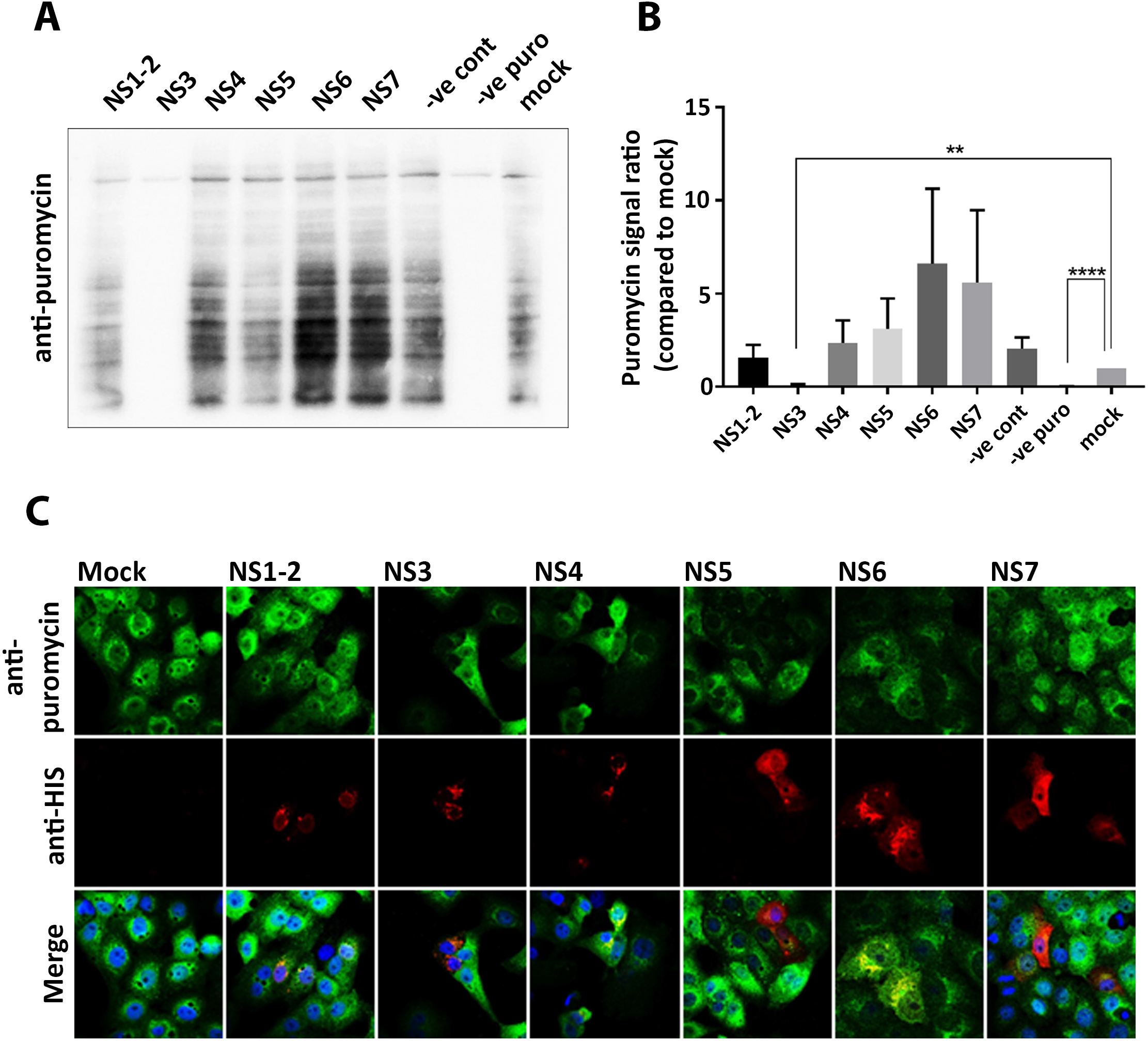
Expression of MNV NS3 protein abolishes host cellular translation: **(A)** HEK 293T cells were transfected with cDNA expression plasmids encoding the individual HIS-tagged MNV NS proteins for 18 hrs, at which point the cells were pulsed with puromycin as previously described and whole cell lysates were obtained and immunolabelled with anti-puromycin antibodies. **(B)** Densitometry analysis of puromycin signal in MNV NS protein transfected cells compared to mock transfected cells (n=3, ANOVA, +/− SEM, **p<0.01, ****p<0.0001). **(C)** Vero cells were transfected with the single HIS-tagged MNV NS proteins and treated with puromycin before cells were fixed and permeabilised for IF. Cells were immunolabelled with antibodies against puromycin (green), 6xHIS (red) and DAPI. Samples were captured via the Zeiss LSM 710 confocal microscope and analysed with the ZEN software.

### MNV-induced suppression of host cell protein translation results in the inability of infected cells to produce major cytokines

The innate immune response is crucial during MNV infection, specifically STAT1 and type I and III IFNs play essential roles in combatting infection (14). We were interested in investigating the impact of impaired protein translation on the innate immune response to MNV infection, and investigated the cytokines IFNβ, TNFα and IL-6 as they are representatives of major immune response pathways. First, we tested whether MNV infection induces the transcriptional activation of IFNβ, TNFα and IL-6. We infected RAW 264.7 with MNV, treated them with Poly(I:C) or left them untreated for 9, 12 and 15 hrs. Transcription levels were assessed using RT-qPCR and compared to mock untreated cells (Fig. 4A). Poly(I:C) stimulation led to the robust induction of IFNβ, TNFα and IL-6 transcription, observed through increasing mRNA levels compared to untreated cells. MNV infected cells also showed similar increases in mRNA levels for both IFNβ and TNFα compared to Poly(I:C) treated cells (Fig 4A, i and ii), however, although we observe a slight increase in IL-6 transcriptional response, this increase was much less profound (Fig. 4A, iii).

To test if the translation of these major cytokines is affected by the global host translation shutoff during MNV infection, we infected RAW 264.7 cells with MNV, treated them with Poly(I:C) and the secretion inhibitor Brefeldin A (BFA), or left them untreated (Fig. 4B). Cell culture supernatant samples were harvested at 9, 12 and 15 h.p.i and cytokine secretion was measured via ELISA. Untreated cells, as well as cells stimulated with Poly(I:C) but treated with BFA, secreted no or only low amounts of IFNβ, TNFα and IL-6 at any time point tested. In contrast, Poly(I:C) stimulated cells released high amounts of all three cytokines into the tissue culture supernatant as early as 9 hrs post treatment. Interestingly, cells infected with MNV showed significantly lower amounts of IFNβ, TNFα and IL-6 being secreted into the cell supernatant compared to Poly(I:C) treated cells (Fig. 4B). Cytokine levels observed for MNV infected cells were similar to Poly(I:C) and BFA treated cells suggesting that secretion might be inhibited, comparable to the function of BFA. Surprisingly, general protein secretion is not disturbed in MNV infected macrophages (Fig. S2), indicating that the reduction in protein levels is likely related to our observed MNV-induced translational inhibition.

**Figure 4.**
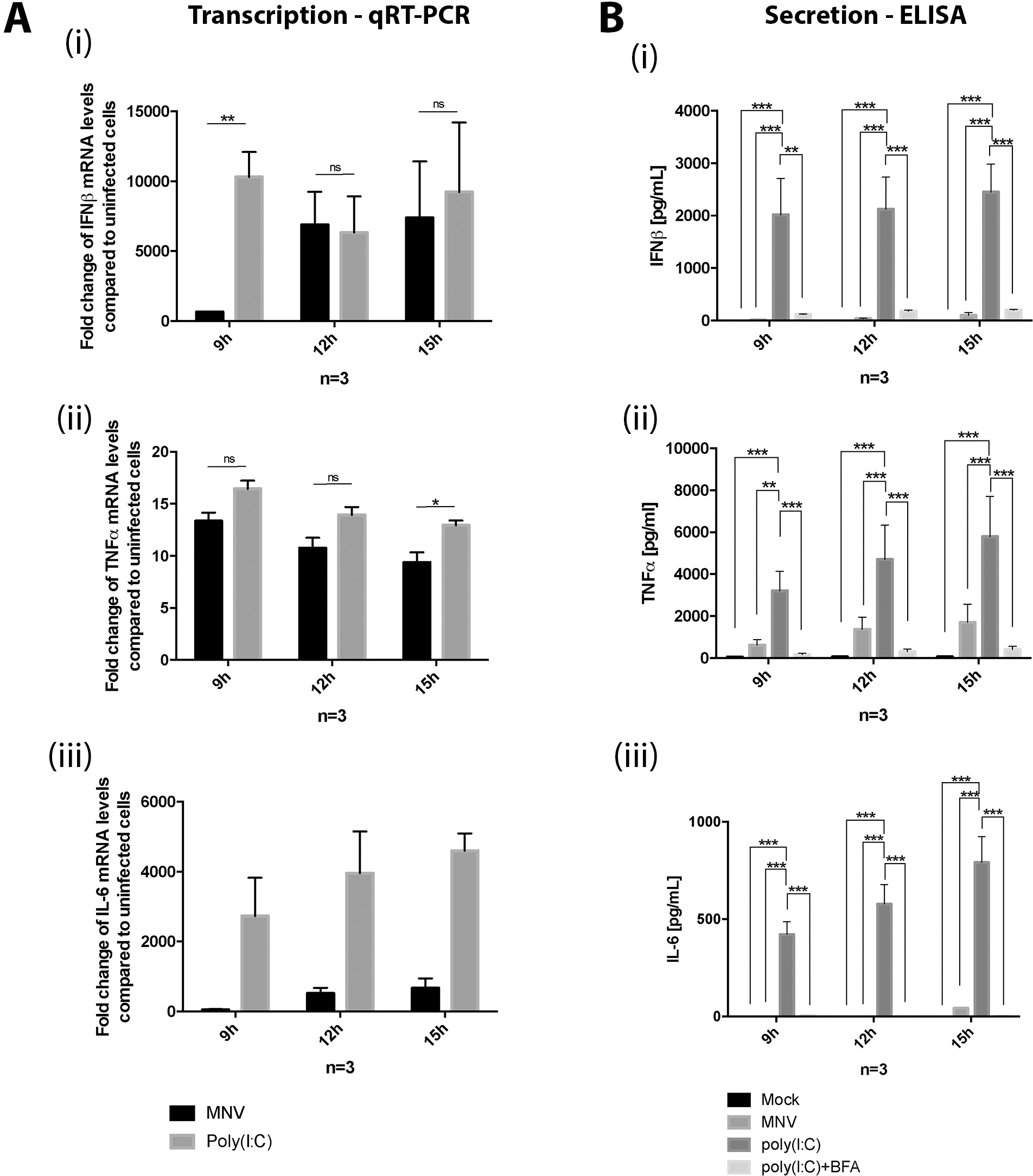
MNV infection in macrophages induces cytokine transcription but inhibits their secretion: **(A)** RAW 264.7 were either MNV-infected (MOI 5) for 9, 12 and 15 hrs or Poly(I:C) treated. RNA samples were taken and analysed via RT-qPCR for the following cytokines (i) IFNβ, (ii) TNFα and (iii) IL-6. **(B)** RAW 264.7 were either mock-infected, infected with MNV (MOI 5) for 9, 12 and 15 hrs, Poly(I:C) or Poly(I:C) + BFA treated. Cell culture supernatants were analysed for the secretion of the specific cytokines (i) IFNβ, (II) TNFα and (III) IL-6 via ELISA. (n=3, ANOVA, +/− SEM, **p<0.01, ****p<0.0001).

### MNV infection inhibits stress granule formation

One of the control mechanisms for translation of interferon stimulated genes (ISGs) and cytokines is the sequestering of the encoding mRNA within cytoplasmic RNA granules *e.g.* SGs (46, 47). Based on our observed profound effect of MNV infection on host cell translation and innate immune associated pathways, we aimed to investigate whether MNV replication also manipulated the formation of SGs. Thus, BMM cells were infected with MNV for 12 hrs and cells were analysed by IF with specific antibodies against the SG marker eIF3η, and the viral VPg protein NS5 (Fig. 5A). In uninfected control cells SG formation was not observed (panels A-C), however, treatment with NaAs induced the formation of numerous and obvious round shaped cytoplasmic puncta (Fig 5A, G-I, arrow head). Interestingly, cells infected with MNV did not appears to contain SGs (panels D-F, arrow head), suggesting that MNV-infection either does not induce SG formation, or that the virus is inhibiting the formation of SGs.

**Figure 5.**
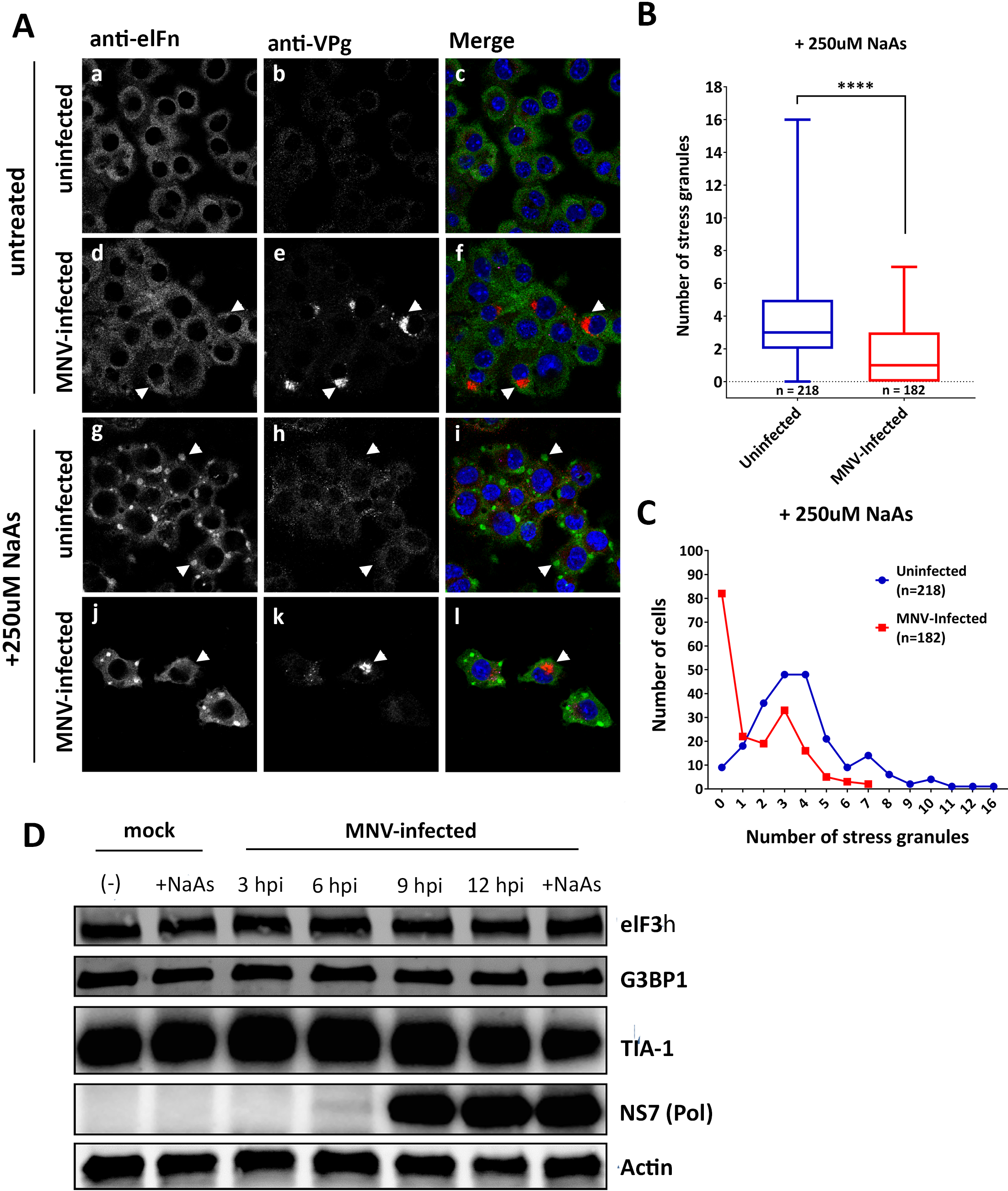
MNV-infection alters SG formation in sodium arsenite treated cells without affecting key SG protein levels. **(A)** BMMs were mock-infected (panels a-c), infected with MNV (MOI 5) for 12 hrs (panels d-f), NaAs treated (250 μM for 20 mins) (panels g-i) or MNV infected and NaAs treated (panels j-l). Cells were fixed for IF analysis and immunolabelled with eIF3η (green), VPg (red, indicating infection) and DAPI (blue). Samples were captured via the Zeiss LSM 710 confocal microscope and analysed with the ZEN software. **(B and C)** SGs in NaAs treated mock (218 cells) and MNV-infected (182 cells) samples were counted from two independent experiments each and collated. **(B)** Box and whiskers plot where whiskers represent min to max and box represents mean with error bars +/− SEM and unpaired two-tailed *t*-test was performed (**** denotes p<0.0001). **(C)** Quantitation demonstrating the total number of cells (y-axis) containing various amount of SGs (x-axis). Blue line represents the number of SGs in uninfected cells (n=218 cells) and the red line represents the number of SGs in MNV-infected cells (n=182 cells) **(D)** BMM cells were either mock-infected or MNV-infected (MOI 5) and whole cell lysates were collected at 3, 6, 9 and 12 hpi. WB was immunolabelled with anti-eIF3η, anti-G3BP1, anti-TIA-1, anti-NS7 and anti-actin antibodies.

To determine if MNV interferes with SG formation, cells were treated with NaAs and subsequently infected with MNV for 12 hrs (Fig. 5A, panels J-L). We observed an inhibitory effect of MNV on the amount of NaAs-induced SGs (Fig. 5A, panels J-L) compared to NaAs-treated uninfected cells (Fig. 5A, panels G-I). In NaAs treated and MNV-infected cells exhibiting SG formation, the morphology of SGs was smaller and elongated, instead of having a typical, round-shaped appearance (Fig. 5A, panels J-L). To quantitate the changes observed, we determined the number of SG foci within MNV infected and uninfected cells in the presence of NaAs (Fig. 5B and C). MNV infected cells displayed a significantly lower number of SGs within the cell compared to uninfected cells during NaAs-treatment. In uninfected cells, an average of 4 SGs per cell were observed (Fig. 5B, blue) with only 4% of cells (9 cells) containing no SGs (Fig. 5C, blue). In contrast, infected cells had only an average of 2 SGs per cell (Fig. 5B, red) and 45% of infected cells (82 cells) contained no SGs at all (Fig. 5C, red). Intriguingly, our observations are contrast to those of Humoud *et al.* (48) who observed that MNV infection did not impact on arsenite-induced SG assembly.

Previous reports have indicated that some viruses prevent the induction of SGs by viral protease-mediated cleavage of G3BP1 and other accessory proteins (49). To determine if this was also true for MNV infection, we investigated the protein levels of eIF3□, G3BP1 and TIA-1 by WB. To this end, we observed no significant change in the total protein levels or size of any of these proteins as infection progressed, indicating that MNV does not manipulate SG formation through protease-mediated cleavage of key SG proteins (Fig 5D). These results suggest that MNV does not induce SGs even though eIF2α is phosphorylated and has an inhibitory effect on SG induction.

### MNV recruits the key SG nucleating proteins G3BP1 to the sites of virus replication which is critical for efficient MNV replication

To examine the ability of MNV to prevent SG induction we visualised the distribution and abundance of key SG nucleating proteins eIF3□ and G3BP1 by IF analysis in MNV-infected BMM cells (Fig. 6). We observed no significant altered distribution of eIF3□ in MNV-infected cells, either at the sites of viral replication or to discrete cytoplasmic foci (Fig. 6A, i compared to a). In contrast, we observed a dramatic redistribution and sequestering of G3BP1 to the sites of MNV replication as identified with antibodies to the MNV VPg protein (NS5) (Fig. 6A, j compared to b). The sequestering of G3BP1 also occurred in MNV-infected cells that were additionally treated with NaAs (Fig. 6A, n compared to f).

**Figure 6.**
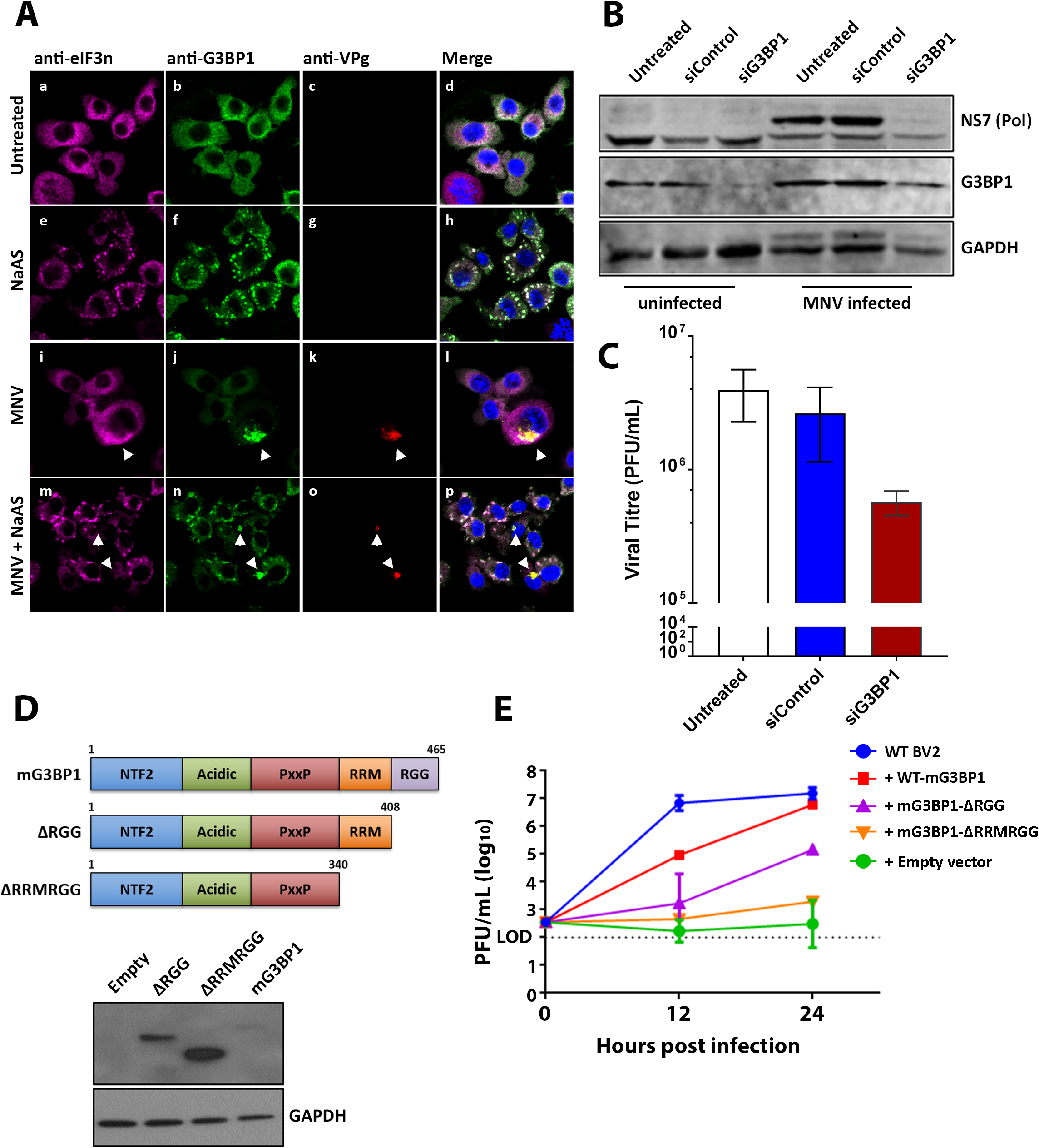
MNV recruits G3BP1 and requires G3BP1 to efficient viral replication. **(A)** BMMs were mock-infected (panels a-d), NaAs treated (250 μM for 20 mins) (panels e-h), infected with MNV (MOI 5) for 12 hrs (panels i-l) or MNV infected and NaAs treated (panels m-p). Cells were fixed for IF analysis and immunolabelled with anti-eIF3η (magenta), anti-G3BP1 (green), VPg (red, indicating infection) and DAPI (blue). Samples were captured via the Zeiss LSM 710 confocal microscope and analysed with the ZEN software. **(B-C)** BMMs were untreated, siControl or siG3BP1 treated and either mock-infected or MNV infected (MOI 5). **(B)** Whole cell lysates were collected for WB (immunolabelled with anti-NS7, anti-G3BP1 and anti-GAPDH) and **(C)** tissue culture fluids were collected for plaque assay. (n=3, mean +/− SEM and unpaired two-tailed t-test was performed). **(D)** Schematic demonstrating WT-mG3BP1, mG3BP1-∆RGG and mG3BP1-∆RRMRGG mutant constructs. Each construct was transfected into BV2 and expression levels were demonstrated by WB. (**E)** WT-BV2 cells, BV2-mG3BP1-KO cells were transfected with WT-mG3BP1, mG3BP1-∆RGG, mG3BP1-∆RRMRGG and an empty vector. Following transfection, cells were infected with MNV and 12 and 24 h.p.i tissue culture fluids were collected for plaque assay. (n=3, mean +/− SEM and unpaired two-tailed t-test was performed).

As we had observed that G3BP1 had been sequestered within the MNV RC, we aimed to determine if this was a functional consequence for evasion of the SG antiviral response or a requirement for replication. Intriguingly in a recent CRISPR screen, G3BP1 was observed as the second most critical host factor, next to the receptor CD300lf, in facilitating MNV infection and replication (50). As the CRISPR knock-out of G3BP1 was observed to be completely inhibit infection, we utilised RNAi-mediated suppression of G3BP1 to identify how MNV may require G3BP1 for replication. Thus, cells were incubated with different RNAi’s specific for the murine *G3bp1* gene and suppression of G3BP1 expression was assessed by WB analysis. siRNA-mediated treatment resulted in a reduction of the G3BP1 protein (Fig. 6B). Upon subsequent infection of these cells we observed an attenuation in MNV replication, by WB (NS7) (Fig. 6B) and the production of infectious virus, as assessed by plaque assays where virus titres with a 1-1/2 logs reduction (representing an ~90-95% decrease in infectious virus) were observed (Fig. 6C).

To further examine the impact of G3BP1 on MNV replication, we knocked out G3BP1 expression in BV2 cells via CRISPR-Cas9 (G3BP1-KO). In G3BP1-KO BV2 cells, we re-introduced G3BP1 by transfecting wild type mouse G3BP1 (WT-mG3BP1) and two mG3BP1 deletion mutants: mG3BP1–∆RGG (∆RGG; deletion of the co-operative RNA binding domain (RGG) from aa408-465) and mG3BP1–∆RRMRGG (∆RRMRGG; additionally includes RNA-binding domain (RRM) from aa340-407) as well as an empty vector (Fig. 6D and E). We infected these cells with MNV and collected virus containing tissue culture fluid at 12 and 24 h.p.i to measure viral titres. In WT-BV2 cells, peak virus replication (10^7^ PFU/mL) was observed after 12 h.p.i and this level remained steady until 24 h.p.i (Fig. 6E, blue line). Consistent with our siRNA results, knockout of G3BP1 resulted in the complete abolishment of virus replication, highlighting the importance of G3BP1 for MNV replication (Fig. 6E, green line). Interestingly, the re-introduction of WT-mG3BP1 into mG3BP1-KO cells completely restored virus replication to WT BV2 levels (10^7^ PFU/mL) by 24 h.p.i, however the rescue of MNV replication was delayed by 12 hrs (Fig. 6E, red line). Furthermore, removal of the G3BP1–∆RGG domain resulted in the partial rescue of MNV replication (10^5^ PFU/mL, 2 logs lower than WT BV2) by 24 h.p.i (Fig. 6E, purple line), however the removal of the G3BP1–∆RRMRGG domains resulted in a 4-log reduction of viral titres (Fig. 6E, orange line)

Based on these observations we suggest that the recruitment of G3BP1 to the MNV RC is essential for virus replication. At this point we have not been able to identify at what stage and how G3BP1contributes to MNV replication, however we would speculate that it contributes to binding of the MNV viral RNA perhaps to stabilise some protein-RNA interactions. It is important to note that sequestration of G3BP1 is also critical to prevent SG formation and promote viral replication without interference of the innate immune response.

## Discussion

The shutdown of host cell translation is one of the major host defence mechanisms against viral infections. Viral replication is completely dependent on host cell translation, as viruses lack their own translational machinery and parasitise the hosts. Therefore, a reduction in host protein translation will likely lead to a decreased translation of viral proteins and interfere with efficient viral replication. Interestingly, translation of viral proteins like NS7, the viral polymerase, does not seem to be affected by the reduced host protein translation during MNV infection, because intracellular amounts of NS7 increase from 6 hpi onwards, while host protein translation subsides (Fig. 1B). These observations strongly suggest that MNV employs an alternative mechanism to initiate translation, independent of cellular protein translation (51). The HuNoV and MNV VPg proteins have been shown to interact with the translation initiation complex through eIF4GI and eIF4E, suggesting a role of NS5 in the initiation of viral protein translation (52, 53). Translation of MNV proteins, which is independent of the cellular cap-dependent protein translation, could be mediated by VPg and allow viral protein translation to occur in the absence of cellular protein translation (54). This would be a great advantage for the virus, not only by forcing the cell to preferentially translate viral proteins, but also by diminishing the innate immune response by preventing the translation of immune effectors such as cytokines.

To uncover how MNV manipulates the ISR, we investigated the PKR/eIF2α pathway, which is a major regulator of the ISR (Fig 1, 2 and 3). We demonstrated that MNV infection leads to the phosphorylation of eIF2α, supporting the observations by Humoud *et al.* (48), and as infection progresses the amount of p-eIF2α drastically increases resulting in timely host cell translational shutoff (Fig. 1). Even though translation was upregulated early during the infection (6 hpi), there was a continuous decrease in the amount of puromycylated proteins at later stages of the infection (from 9 hpi), indicating a reduction in global host cell translation (Fig. 1B). Based on the immunoblot and IF analysis (Fig. 1C), MNV starts to affect host cell translation from 9 hpi, reducing host cell translation to a minimum in most infected cells by 12 hpi (Fig. 1C). Expression studies of single viral proteins revealed that NS3 expression alone is sufficient to induce translation inhibition (Fig. 3).

We presumed that p-eIF2α may be regulated via PKR which is activated by binding to dsRNA produced during MNV infection. We initially showed that phosphorylation of eIF2α was mediated via PKR rather than PERK (Fig 2). Interestingly though, when cells were treated with C16 and analysed for their translation activity via puromycin treatment, we did not observe an increase in host cell translation activity in MNV-infected cells compared to infected, but untreated cells (Fig 2B). These observations show that the PKR/eIF2α axis is activated during MNV infection but is not solely responsible for the host cell translation shutdown. However, the shutdown of host translation by MNV is effective and robust (Fig. 3) and we show that it affects the translation of innate immune response regulators like cytokines (Fig. 4).

The release of cytokines such as IFNβ, TNFα and IL-6 during viral infections plays a crucial role in the innate immune response against viruses. The transcription and translation of cytokines are elevated in virus infected cells, mostly due to the recognition of PAMPs, e.g. dsRNA. Cytokines are then secreted into the extracellular space where they can either bind to receptors on neighbouring cells or to receptors on the infected cell itself to enhance the antiviral response (55). We and others have shown that MNV infected cells increase the transcription of cytokine (IFNβ, TNFα and IL-6) mRNAs (Fig. 4) (56, 57) indicating that PRRs like MDA5 (58) have successfully detected the viral infection and activated an antiviral response against it. Intriguingly, MNV infected cells do not secrete significant levels cytokines which would help to overcome and contain the acute infection (Fig. 4). Our subsequent studies revealed that the low amounts of secreted cytokines from infected cells were not due to the inhibition of general protein secretion (Fig. S2). Instead, we observed that only very low amounts of translated cytokines can be detected within the infected cells, further confirming that there is no secretion inhibition, which would cause the accumulation of cytokines within the cells (Fig. 4). The difference in intracellular protein levels for TNFα compared to the mRNA levels indicates an interference of the virus with either protein stability or the translation of host cell proteins.

Like all viruses, MNV must modulate host responses to provide conditions suitable for intracellular replication. To replicate successfully, MNV must also control the localisation of viral RNA within the host cell, as these replication by-products are highly immuno-stimulatory. SG formation, which is part of the ISR, generates cytoplasmic granules containing stalled translational machineries involved in regulating RNA transcript homeostasis (59). This mechanism serves as an extension of translation by sequestering mRNA from active translation, whilst allowing the translation of certain mRNAs. This translational regulation is typically induced upon exposure to cellular stresses including ER stress, oxidative stress, heat shock (60, 61) and viral infection (59). Under stressed conditions, cells activate eIF2α kinases to phosphorylate eIF2α which depletes the eIF2α-GTP-tRNA^met^ ternary complex required to form the preinitiation complex, resulting in stalled translation initiation (31, 32). These stalled preinitiation complexes aggregate and form SGs, thereby general protein translation is inhibited (30). This host response to infection can affect cytokine translation, thus some viruses have devised strategies to regulate RNA granule function to selectively control RNA translation, and therefore promote their replication (Reviewed in 28, 62).

Previous studies have shown that many different virus families modulate SG function to allow efficient replication (63). Thus, as we observed eIF2a phosphorylation in MNV infected cells, we extended our IF analysis to examine whether SGs form during MNV infection. We demonstrated via IF that SGs are reduced during MNV infection, although eIF2α is phosphorylated. In fact, when cells are treated with NaAs, MNV infection significantly dampened SG formation and SGs that were present had atypical morphologies and reduced SG numbers (Fig. 5A and B). These results not only suggest MNV infection does not induce SG formation, but MNV can also exert control over SG formation. Intriguingly, Humoud *et al*. (48) did not observe these findings and it is difficult to reconcile their findings to ours. The only difference is they utilised J774 macrophages in their study potentially identifying subtle cell type differences.

We have shown that MNV recruits G3BP1 to sites of viral replication (Fig. 6A) and demonstrate through siRNA-mediated knockdown and CRISPR-Cas9 depletion of G3BP1 that it is required for efficient viral replication (Figs. 6). We postulate that MNV recruits G3BP1 to sites of viral replication where G3BP1 binds to viral RNA via the RNA Recognising Motif (RRM). G3BP1 recruitment to the assembly complex seems to serve a dual purpose, i) the promotion of viral replication (presumably by aiding in RNA duplex unwinding), and ii) the prevention of SG formation.

Based on our findings, MNV likely employs a strategy similar to picornaviruses and alphaviruses to evade the innate immune response by inducing the inhibition of host cell translation (64–68). During MNV infection, shutdown of host translation is independent of the SG-PKR-eIF2α axis and PABP cleavage and seems to be regulated through an unknown mechanism (Fig. 7, model). It will be interesting to investigate if MNV cleaves other components of the translation complex, or if it regulates the host translation shutdown through another pathway, like miRNA or preferred binding of viral mRNA to the translation complex. Overall, it is important to note that the MNV NS3 protein allows the inhibition of cap-dependent host cell translation, while inhibiting the formation of SGs by recruiting G3BP1 to the MNV RC, which sequester stalled pre-translation complexes containing essential components of the translational machinery (Fig. 7, model). It is intriguing to speculate that MNV selectively induces cap-dependent translation inhibition to enhance viral translation and inhibit the innate immune response but needs access to the translational machinery and therefore inhibits SG formation. This strategy not only increases viral replication efficiency, but also promotes immune evasion of the virus, which could explain the rapid replication cycle and the delayed innate immune response to MNV infection.

**Figure 7.**
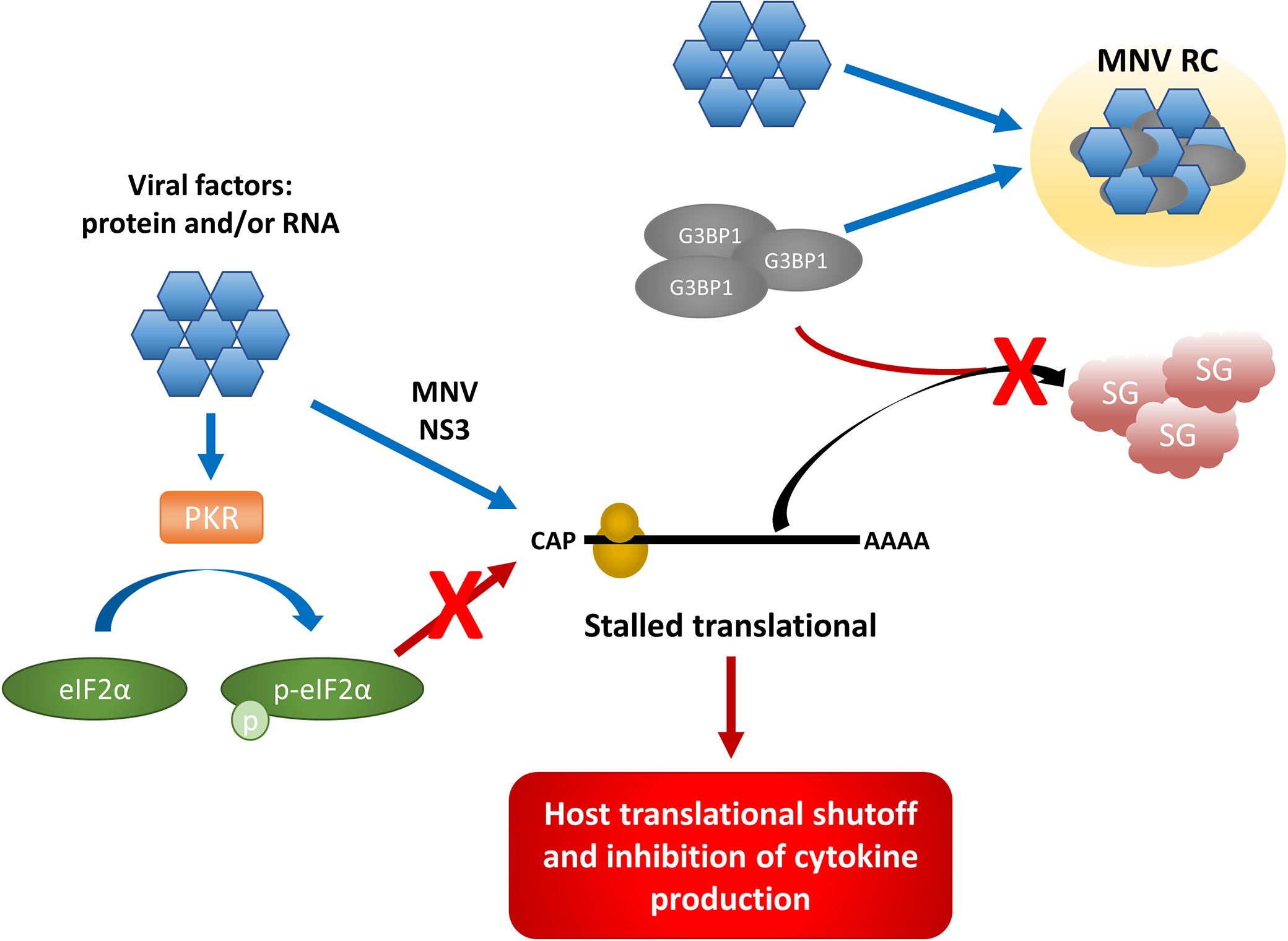
Model. During MNV infection, viral factors such as proteins and/or RNA (blue hexagon) phosphorylate eIF2α (green oval) via PKR (orange rectangle), as well as stalling translation initiation by the MNV NS3 protein, however this translational arrest is uncoupled from the PKR–p-eIF2α axis. These stalled preinitiation complexes typically aggregate with G3BP1 (grey oval) and form SGs (red cloud), however MNV viral factors sequester G3BP1 to the MNV RC (yellow circle) to promote replication. This allows the inhibition of cap-dependent host cell translation, as well as inhibiting the formation of SGs.

## Materials and Methods

### Cell lines

RAW 264.7 murine macrophages, bone marrow-derived macrophages (BMM) Vero and HEK 293T cells were maintained in Dulbecco’s Modified Eagle’s Medium (DMEM) (Gibco) supplemented with 10% foetal calf serum (FCS) (Gibco) and 1% GlutaMAX (200mM) (Gibco). BV2 cells were cultured in DMEM with 10% FBS, 1% HEPES and 1% GlutaMAX. All cell lines were cultivated at 37°C in a 5% CO_2_ incubator, as previously described (19).

### MNV infection

RAW 264.7 macrophages, BMMs and BV2 cells were infected with MNV at a multiplicity of infection (MOI) of 5, as previously described (19). Cells were rocked in a low volume of media for one hour at 37 °C, before cells were supplemented with additional media. Unless indicated differently, cells were fixed or lysed at 12 h.p.i. If supernatant was collected, the media was centrifuged at 10,000g for 3 min to pellet cellular debris.

### Chemicals and Antibodies

Sodium arsenite (Sigma-Aldrich) was added to cells at a concentration of 250 μM for 20 mins prior to fixation or cell lysate collection. The PKR inhibitor C16 (Sigma-Aldrich) was added to infected cells at a concentration of 1 μM at 1 h.p.i and cell lysates were collected 12 h.p.i. The ISR inhibitor ISRIB (Sigma-Aldrich) was added 1 h.p.i at 0.5 μM. Puromycin (Life Technologies) was added to cells at a concentration of 10 μg/ml at indicated times prior to cell lysate collection. Goat anti-eIF3η, Goat anti-G3BP1 and Goat anti-TIA-1 were all purchased from Santa Cruz Biotech. Rabbit anti-eIF2α was purchased from Invitrogen; Rabbit anti-actin from Sigma-Aldrich; Mouse anti-puromycin from Kerafast Inc; Mouse anti-G3BP1, Mouse anti-GAPDH, Rabbit anti-HIS and Rabbit anti-calnexin from Abcam and Rabbit anti-p-eIF2α (S52) and Alexa Fluor-conjugated species-specific IgG were purchased from Life Technologies. Rabbit anti-NS7 and Rabbit anti-NS5 were manufactured and produced by Invitrogen.

### Plaque Assay

3.0 x 10^5^ RAW264.7 or BV2 cells were seeded onto 12 well plates and incubated 37°C until 70% confluent. Virus-containing supernatants were ten-fold serially diluted in DMEM, added to plates and rocked every 10 mins for 1 hr at 37°C. Following incubation, plaque assay overlay (70% DMEM, 2.5% v/v FCS, 13.3 mM NaHCO_3_, 22.4 mM 4-(2-hydroxyethyl)-1-piperazineethanesulfonic acid (HEPES), 200 mM GlutaMAX and 0.35 % w/v low-melting-point (LMP) agarose) was added to each well. Overlay was solidified at 4°C for 15 mins and incubated at 37°C for 48 hrs. Cells were fixed in 10% formalin for 1 hr at RT. Plaque assay overlay was removed and cells were stained with 1 ml of Toludine blue for 30 min. Stain was removed, rinsed with water and plaque formations were enumerated.

### Immunofluorescence microscopy

Cells were rinsed twice with Phosphate buffered saline (PBS) and fixed 4% v/v paraformaldehyde (PFA)/PBS for 15 min at RT. Fixative was removed and cells were permeabilised with 0.1% v/v Triton X-100 for 10 min at RT. Cells were rinsed twice with PBS and quenched with 0.2 M glycine for 10 mins at RT. Cells were then rinsed with PBS and coverslips were incubated in primary antibodies diluted in 25 μl of 1% bovine serum albumin (BSA)/PBS for 1 hr at RT. Following incubation with primary antibodies, cells were washed thrice with 0.1% BSA/PBS. Coverslips were incubated in secondary antibodies diluted in 25 μl of 1 % BSA/PBS for 45 min at RT. Cells were washed twice with PBS and incubated for 5 mins with 4,6-diamidino-2-phenylindole (DAPI) (0.33 μg/ml) in PBS. Coverslips were rinsed twice with PBS and MilliQ water and mounted on cover-slides with ProLong Diamond (Life Technologies). Cells were analysed using the Zeiss LSM710 confocal microscope.

### Western Blot Analysis

Cells were lysed with NP-40 lysis buffer containing protease inhibitor cocktail and phosphatase inhibits. Samples were separated on a bis/tris polyacrylamide gel and transferred to a PVDF membrane. The membrane was blocked with 5% BSA/PBS-T for 2 h. Primary antibodies were added in 5 % BSA/PBS-T and incubated overnight at 4 **°**C. The following day, the membrane was washed 3 x with PBS-T and then incubated with secondary antibody in PBS-T for 90 mins at RT. The membrane was then washed four times in PBS-T then visualised using the MF-ChemiBIS DNR (Bio-imaging Systems).

### ELISA

Cell culture supernatants were analysed for their cytokine concentration using mouse specific

ELISA kits for the following cytokines: IFNβ (LEGEND MAXTM, BioLegend) and TNFα (ELISAkit.com). Tissue culture supernatants and standards were applied to the 96-well pre-coated assay plate and incubated for 2 h. Wells were washed 4 x in assay wash buffer, before adding the assay specific biotin-labelled detection antibody. Plates were incubated for 2 hrs at room temperature, wells were washed again 4 x with the assay wash buffer. A streptavidin conjugated HRP was added afterwards and incubated for a further 45 min. Plates were washed thoroughly 6 x with wash buffer and the TMB substrate was added. Plates were checked every 3-5 min to observe the colour change. The reaction was stopped using the assay stop solution. Absorbance at 450 nm and 570 nm (background) was measured using the CLARIOstar® microplate. Cytokine concentrations were calculated using the ELISAanalysis.com website.

### qRT-PCR

Cells for RNA extraction were lysed with Trizol (Life Technologies). The total RNA was isolated by phenol:chloroform extraction and then stored at −80°C. Total RNA concentration was quantified using a Nanodrop and 1 μg of total RNA was treated with RQ1 DNase (Promega) at 37°C for 45 mins. cDNA was generated by reverse-transcription using Sensifast RT (Bioline) at 25°C for 10 mins, 42°C for 15 mins and 85°C for 5 mins. cDNA levels were quantified by qPCR with Sybr GreenER (Bio-Rad) using the following cycling conditions (50°C for 8 mins, 95°C for 2 mins, 40 cycles of 15 secs at 95°C, 1 min annealing/extension at 60°C followed by final extension of 10 mins). Fold induction of RNA was compared to the housekeeping gene (GADPH) and error bars indicate mean +/− SEM from triplicate experiments.

### RNAi-mediated depletion of G3BP1

BMM cells were reverse transfected with 0.25 μM siRNA (Bioneer) and RNAiMAX (Invitrogen) in opti-MEM (Gibco). Cells were incubated at 37°C, 5% CO2 for 24 hrs. The following day, cells were once again transfected with 0.5 μM siRNA. 24 hrs post transfection; cells were infected with MNV at a MOI of 5 for 1 hour, transfected with 0.5 μM siRNA and incubated for a further 12 hrs. Twelve hrs post infection, whole cell lysates and supernatants were collected for WB and plaque assay, respectively.

### Generation of G3BP1 KO cells via CRISPR-Cas9

BV2 cells were cultured in DMEM containing 10% FBS and 1% HEPES. BV2 cells were transiently transfected with Cas9 and a sgRNA (5’ TTCCCCGGCCCCGGCTGATGNGG 3’) targeting exon 7 of G3BP1. BV2 cells were then single cell cloned and G3BP1 was sequenced by Illumina HiSeq. BV2 cells are polyploid at the G3BP1 locus as described previously (PMID: 27540007). Clone 1A3 had four independent deletions at the sgRNA binding site resulting in two unique 1 base pair deletions in addition to 4 and 10 base pair deletions. The mutations introduced resulted in frame shifts and the absence of detectable G3BP1 protein as measured by WB. Sequences are available upon request.

## Authors Contributions

### Conceptualization

Jason M. Mackenzie, Craig B. Wilen, Peter A. White

### Data curation

Turgut E. Aktepe, Svenja Fritzlar, Yi-Wei Chao, Michael R. McAllaster

### Formal analysis

Turgut E. Aktepe, Svenja Fritzlar, Yi-Wei Chao, Michael R. McAllaster

### Funding acquisition

Jason M. Mackenzie, Craig B. Wilen, Peter A. White

### Investigation

Turgut E. Aktepe, Svenja Fritzlar, Yi-Wei Chao, Michael R. McAllaster

### Methodology

Turgut E. Aktepe, Svenja Fritzlar, Yi-Wei Chao, Michael R. McAllaster, Craig B. Wilen, Peter A. White, Jason M. Mackenzie

### Project administration

Jason M. Mackenzie, Craig B. Wilen, Peter A. White

### Supervision

Jason M. Mackenzie, Craig B. Wilen, Peter A. White

### Validation

Turgut E. Aktepe, Svenja Fritzlar, Yi-Wei Chao, Michael R. McAllaster, Craig B. Wilen, Jason M. Mackenzie

### Visualization

Turgut E. Aktepe, Svenja Fritzlar, Jason M. Mackenzie

### Writing – original draft preparation

Turgut E. Aktepe, Svenja Fritzlar, Jason M. Mackenzie

### Writing – review and editing

Turgut E. Aktepe, Svenja Fritzlar, Yi-Wei Chao, Michael R. McAllaster, Craig B. Wilen, Peter A. White, Jason M. Mackenzie

## Supporting information

Supplementary Figue 1

Supplementary Figue 2

## Acknowledgements

We thank Herbert “Skip” Virgin for his help, knowledge and guidance throughout this study. This work was funded by a National Health and Medical Research Council project grants (APP1083139 and APP1123135) awarded to JMM and PAW. CBW was supported by NIH grant K08 A128043.

## References

1. Hall AJ, Lopman BA, Payne DC, Patel MM, Gastanaduy PA, Vinje J, Parashar UD. 2013. Norovirus disease in the United States. Emerg Infect Dis 19:1198–205.

2. Lopman BA, Hall AJ, Curns AT, Parashar UD. 2011. Increasing rates of gastroenteritis hospital discharges in US adults and the contribution of norovirus, 1996-2007. Clin Infect Dis 52:466–74.

3. Scallan E, Hoekstra RM, Angulo FJ, Tauxe RV, Widdowson MA, Roy SL, Jones JL, Griffin PM. 2011. Foodborne illness acquired in the United States–major pathogens. Emerg Infect Dis 17:7–15.

4. Adler JL, Zickl R. 1969. Winter vomiting disease. J Infect Dis 119:668–73.

5. Rockx B, De Wit M, Vennema H, Vinje J, De Bruin E, Van Duynhoven Y, Koopmans M. 2002. Natural history of human calicivirus infection: a prospective cohort study. Clin Infect Dis 35:246–53.

6. Graham DY, Jiang X, Tanaka T, Opekun AR, Madore HP, Estes MK. 1994. Norwalk virus infection of volunteers: new insights based on improved assays. J Infect Dis 170:34–43.

7. Atmar RL, Bernstein DI, Harro CD, Al-Ibrahim MS, Chen WH, Ferreira J, Estes MK, Graham DY, Opekun AR, Richardson C, Mendelman PM. 2011. Norovirus vaccine against experimental human Norwalk Virus illness. N Engl J Med 365:2178–87.

8. Atmar RL, Estes MK. 2012. Norovirus vaccine development: next steps. Expert Rev Vaccines 11:1023–5.

9. Lindesmith LC, Mallory ML, Jones TA, Richardson C, Goodwin RR, Baehner F, Mendelman PM, Bargatze RF, Baric RS. 2017. Impact of Pre-exposure History and Host Genetics on Antibody Avidity Following Norovirus Vaccination. J Infect Dis 215:984–991.

10. Lindesmith LC, Ferris MT, Mullan CW, Ferreira J, Debbink K, Swanstrom J, Richardson C, Goodwin RR, Baehner F, Mendelman PM, Bargatze RF, Baric RS. 2015. Broad blockade antibody responses in human volunteers after immunization with a multivalent norovirus VLP candidate vaccine: immunological analyses from a phase I clinical trial. PLoS Med 12:e1001807.

11. El-Kamary SS, Pasetti MF, Mendelman PM, Frey SE, Bernstein DI, Treanor JJ, Ferreira J, Chen WH, Sublett R, Richardson C, Bargatze RF, Sztein MB, Tacket CO. 2010. Adjuvanted intranasal Norwalk virus-like particle vaccine elicits antibodies and antibody-secreting cells that express homing receptors for mucosal and peripheral lymphoid tissues. J Infect Dis 202:1649–58.

12. Jones MK, Watanabe M, Zhu S, Graves CL, Keyes LR, Grau KR, Gonzalez-Hernandez MB, Iovine NM, Wobus CE, Vinje J, Tibbetts SA, Wallet SM, Karst SM. 2014. Enteric bacteria promote human and mouse norovirus infection of B cells. Science 346:755–9.

13. Ettayebi K, Crawford SE, Murakami K, Broughman JR, Karandikar U, Tenge VR, Neill FH, Blutt SE, Zeng XL, Qu L, Kou B, Opekun AR, Burrin D, Graham DY, Ramani S, Atmar RL, Estes MK. 2016. Replication of human noroviruses in stem cell-derived human enteroids. Science 353:1387–1393.

14. Wobus CE, Karst SM, Thackray LB, Chang K-O, Sosnovtsev SV, Belliot G, Krug A, Mackenzie JM, Green KY, Virgin IV HW. 2004. Replication of Norovirus in cell culture reveals a tropism for dendritic cells and macrophages. PLoS Biol 2:e432.

15. Karst SM, Wobus CE, Lay M, Davidson J, Virgin HWt. 2003. STAT1-dependent innate immunity to a Norwalk-like virus. Science 299:1575–8.

16. McFadden N, Bailey D, Carrara G, Benson A, Chaudhry Y, Shortland A, Heeney J, Yarovinsky F, Simmonds P, Macdonald A, Goodfellow I. 2011. Norovirus regulation of the innate immune response and apoptosis occurs via the product of the alternative open reading frame 4. PLoS Pathog 7:e1002413.

17. Daughenbaugh KF, Wobus CE, Hardy ME. 2006. VPg of murine norovirus binds translation initiation factors in infected cells. Virol J 3:33.

18. Goodfellow I, Chaudhry Y, Gioldasi I, Gerondopoulos A, Natoni A, Labrie L, Laliberte JF, Roberts L. 2005. Calicivirus translation initiation requires an interaction between VPg and eIF 4 E. EMBO Rep 6:968–72.

19. Hyde JL, Sosnovtsev SV, Green KY, Wobus C, Virgin HW, Mackenzie JM. 2009. Mouse norovirus replication is associated with virus-induced vesicle clusters originating from membranes derived from the secretory pathway. J Virol 83:9709–19.

20. Hyde JL, Gillespie LK, Mackenzie JM. 2012. Mouse Norovirus 1 Utilizes the Cytoskeleton Network To Establish Localization of the Replication Complex Proximal to the Microtubule Organizing Center. Journal of Virology 86:4110.

21. McCune BT, Tang W, Lu J, Eaglesham JB, Thorne L, Mayer AE, Condiff E, Nice TJ, Goodfellow I, Krezel AM, Virgin HW. 2017. Noroviruses Co-opt the Function of Host Proteins VAPA and VAPB for Replication via a Phenylalanine–Phenylalanine-Acidic-Tract-Motif Mimic in Nonstructural Viral Protein NS1/2. mBio 8:e00668–17.

22. Hyde JL, Mackenzie JM. 2010. Subcellular localization of the MNV-1 ORF1 proteins and their potential roles in the formation of the MNV-1 replication complex. Virology 406:138–48.

23. Cotton BT, Hyde JL, Sarvestani ST, Sosnovtsev SV, Green KY, White PA, Mackenzie JM. 2017. The Norovirus NS3 Protein Is a Dynamic Lipid- and Microtubule-Associated Protein Involved in Viral RNA Replication. J Virol 91.

24. Zamyatkin DF, Parra F, Alonso JMM, Harki DA, Peterson BR, Grochulski P, Ng KK-S. 2008. Structural insights into mechanisms of catalysis and inhibition in Norwalk virus polymerase. Journal of Biological Chemistry 283:7705–7712.

25. Högbom M, Jäger K, Robel I, Unge T, Rohayem J. 2009. The active form of the norovirus RNA-dependent RNA polymerase is a homodimer with cooperative activity. Journal of General Virology 90:281–291.

26. Leen EN, Baeza G, Curry S. 2012. Structure of a murine norovirus NS6 protease-product complex revealed by adventitious crystallisation. PloS one 7:e38723.

27. Fernandes H, Leen EN, Cromwell Jr H, Pfeil M-P, Curry S. 2015. Structure determination of Murine Norovirus NS6 proteases with C-terminal extensions designed to probe protease-substrate interactions. PeerJ 3:e798.

28. Lloyd RE. 2013. Regulation of stress granules and P-bodies during RNA virus infection. Wiley Interdiscip Rev RNA 4:317–31.

29. Karst SM, Wobus CE, Lay M, Davidson J, Virgin HW, IV. 2003. STAT1-Dependent Innate Immunity to a Norwalk-Like Virus. Science 299:1575–1578.

30. Kedersha NL, Gupta M, Li W, Miller I, Anderson P. 1999. RNA-binding proteins TIA-1 and TIAR link the phosphorylation of eIF-2 alpha to the assembly of mammalian stress granules. J Cell Biol 147:1431–42.

31. Duncan RF, Hershey JW. 1987. Translational repression by chemical inducers of the stress response occurs by different pathways. Arch Biochem Biophys 256:651–61.

32. McEwen E, Kedersha N, Song B, Scheuner D, Gilks N, Han A, Chen JJ, Anderson P, Kaufman RJ. 2005. Heme-regulated inhibitor kinase-mediated phosphorylation of eukaryotic translation initiation factor 2 inhibits translation, induces stress granule formation, and mediates survival upon arsenite exposure. J Biol Chem 280:16925–33.

33. Kedersha N, Chen S, Gilks N, Li W, Miller IJ, Stahl J, Anderson P. 2002. Evidence that ternary complex (eIF2-GTP-tRNAi Met)–deficient preinitiation complexes are core constituents of mammalian stress granules. Molecular biology of the cell 13:195–210.

34. Gorchakov R, Frolova E, Williams BR, Rice CM, Frolov I. 2004. PKR-dependent and-independent mechanisms are involved in translational shutoff during Sindbis virus infection. Journal of virology 78:8455–8467.

35. Khaperskyy DA, Hatchette TF, McCormick C. 2012. Influenza A virus inhibits cytoplasmic stress granule formation. FASEB J 26:1629–39.

36. White JP, Cardenas AM, Marissen WE, Lloyd RE. 2007. Inhibition of cytoplasmic mRNA stress granule formation by a viral proteinase. Cell Host Microbe 2:295–305.

37. Esclatine A, Taddeo B, Roizman B. 2004. Herpes simplex virus 1 induces cytoplasmic accumulation of TIA-1/TIAR and both synthesis and cytoplasmic accumulation of tristetraprolin, two cellular proteins that bind and destabilize AU-rich RNAs. J Virol 78:8582–92.

38. Emara MM, Brinton MA. 2007. Interaction of TIA-1/TIAR with West Nile and dengue virus products in infected cells interferes with stress granule formation and processing body assembly. Proc Natl Acad Sci U S A 104:9041–6.

39. Clemens MJ. 2001. Initiation factor eIF2 alpha phosphorylation in stress responses and apoptosis. Prog Mol Subcell Biol 27:57–89.

40. McInerney GM, Kedersha NL, Kaufman RJ, Anderson P, Liljestrom P. 2005. Importance of eIF2alpha phosphorylation and stress granule assembly in alphavirus translation regulation. Mol Biol Cell 16:3753–63.

41. Balachandran S, Roberts PC, Brown LE, Truong H, Pattnaik AK, Archer DR, Barber GN. 2000. Essential role for the dsRNA-dependent protein kinase PKR in innate immunity to viral infection. Immunity 13:129–141.

42. He B. 2006. Viruses, endoplasmic reticulum stress, and interferon responses. Cell death and differentiation 13:393.

43. Shimazawa M, Hara H. 2006. Inhibitor of double stranded RNA-dependent protein kinase protects against cell damage induced by ER stress. Neuroscience Letters 409:192–195.

44. Sidrauski C, Acosta-Alvear D, Khoutorsky A, Vedantham P, Hearn BR, Li H, Gamache K, Gallagher CM, Ang KKH, Wilson C, Okreglak V, Ashkenazi A, Hann B, Nader K, Arkin MR, Renslo AR, Sonenberg N, Walter P. 2013. Pharmacological brake-release of mRNA translation enhances cognitive memory. eLife 2:e00498.

45. Emmott E, Sorgeloos F, Caddy SL, Vashist S, Sosnovtsev S, Lloyd R, Heesom K, Locker N, Goodfellow I. 2017. Norovirus-Mediated Modification of the Translational Landscape via Virus and Host-Induced Cleavage of Translation Initiation Factors. Molecular & Cellular Proteomics 16:S215.

46. Li MMH, MacDonald MR, Rice CM. 2015. To translate, or not to translate: viral and host mRNA regulation by interferon-stimulated genes. Trends in Cell Biology 25:320–329.

47. Anderson P. 2008. Post-transcriptional control of cytokine production. Nature Immunology 9:353.

48. Humoud MN, Doyle N, Royall E, Willcocks MM, Sorgeloos F, van Kuppeveld F, Roberts LO, Goodfellow IG, Langereis MA, Locker N. 2016. Feline Calicivirus Infection Disrupts Assembly of Cytoplasmic Stress Granules and Induces G3BP1 Cleavage. Journal of Virology 90:6489.

49. Reineke LC, Lloyd RE. 2013. Diversion of stress granules and P-bodies during viral infection. Virology 436:255–267.

50. Orchard RC, Wilen CB, Doench JG, Baldridge MT, McCune BT, Lee YC, Lee S, Pruett-Miller SM, Nelson CA, Fremont DH, Virgin HW. 2016. Discovery of a proteinaceous cellular receptor for a norovirus. Science 353:933–6.

51. Hanson PJ, Zhang HM, Hemida MG, Ye X, Qiu Y, Yang D. 2012. IRES-Dependent Translational Control during Virus-Induced Endoplasmic Reticulum Stress and Apoptosis. Frontiers in Microbiology 3:92.

52. Daughenbaugh KF, Fraser CS, Hershey JW, Hardy ME. 2003. The genome‐linked protein VPg of the Norwalk virus binds eIF3, suggesting its role in translation initiation complex recruitment. The EMBO journal 22:2852–2859.

53. Daughenbaugh KF, Wobus CE, Hardy ME. 2006. VPg of murine norovirus binds translation initiation factors in infected cells. Virology journal 3:1.

54. Chung L, Bailey D, Leen EN, Emmott EP, Chaudhry Y, Roberts LO, Curry S, Locker N, Goodfellow IG. 2014. Norovirus translation requires an interaction between the C terminus of the genome-linked viral protein VPg and eukaryotic translation initiation factor 4G. Journal of Biological Chemistry 289:21738–21750.

55. Martinez FO, Sica A, Mantovani A, Locati M. 2008. Macrophage activation and polarization. Front Biosci 13:453–461.

56. Enosi Tuipulotu D, Netzler NE, Lun JH, Mackenzie JM, White PA. 2017. RNA Sequencing of Murine Norovirus-Infected Cells Reveals Transcriptional Alteration of Genes Important to Viral Recognition and Antigen Presentation. Front Immunol 8:959.

57. Levenson EA, Martens C, Kanakabandi K, Turner CV, Virtaneva K, Paneru M, Ricklefs S, Sosnovtsev SV, Johnson JA, Porcella SF, Green KY. 2018. Comparative Transcriptomic Response of Primary and Immortalized Macrophages to Murine Norovirus Infection. J Immunol 200:4157–4169.

58. McCartney SA, Thackray LB, Gitlin L, Gilfillan S, Virgin HW, Colonna M. 2008. MDA-5 recognition of a murine norovirus. PLoS Pathog 4:e1000108.

59. Mazroui R, Sukarieh R, Bordeleau ME, Kaufman RJ, Northcote P, Tanaka J, Gallouzi I, Pelletier J. 2006. Inhibition of ribosome recruitment induces stress granule formation independently of eukaryotic initiation factor 2alpha phosphorylation. Mol Biol Cell 17:4212–9.

60. Anderson P, Kedersha N. 2006. RNA granules. J Cell Biol 172:803–8.

61. Holcik M, Sonenberg N. 2005. Translational control in stress and apoptosis. Nat Rev Mol Cell Biol 6:318–27.

62. Khong A, Jan E. 2011. Modulation of stress granules and P bodies during dicistrovirus infection. J Virol 85:1439–51.

63. Lindquist ME, Lifland AW, Utley TJ, Santangelo PJ, Crowe JE, Jr. 2010. Respiratory syncytial virus induces host RNA stress granules to facilitate viral replication. J Virol 84:12274–84.

64. Mosenkis J, Daniels-McQueen S, Janovec S, Duncan R, Hershey JW, Grifo JA, Merrick WC, Thach RE. 1985. Shutoff of host translation by encephalomyocarditis virus infection does not involve cleavage of the eucaryotic initiation factor 4F polypeptide that accompanies poliovirus infection. J Virol 54:643–5.

65. Etchison D, Milburn S, Edery I, Sonenberg N, Hershey J. 1982. Inhibition of HeLa cell protein synthesis following poliovirus infection correlates with the proteolysis of a 220,000-dalton polypeptide associated with eucaryotic initiation factor 3 and a cap binding protein complex. Journal of Biological Chemistry 257:14806–14810.

66. Lloyd R, Grubman M, Ehrenfeld E. 1988. Relationship of p220 cleavage during picornavirus infection to 2A proteinase sequencing. Journal of virology 62:4216–4223.

67. Lloyd RE, Jense H, Ehrenfeld E. 1987. Restriction of translation of capped mRNA in vitro as a model for poliovirus-induced inhibition of host cell protein synthesis: relationship to p220 cleavage. Journal of virology 61:2480–2488.

68. McInerney GM, Kedersha NL, Kaufman RJ, Anderson P, Liljestrom P. 2005. Importance of eIF2α phosphorylation and stress granule assembly in alphavirus translation regulation. Molecular biology of the cell 16:3753–3763.

