## Supplementary data text file for "Mouse Norovirus infection arrests host cell translation uncoupled from the stress granule-PKR-eIF2α axis"

^1^ Department of Microbiology and Immunology, University of Melbourne at the Peter Doherty Institute for Infection and Immunity, Melbourne, VIC, Australia, ^2^ Department of Pathology and Immunology, Washington University School of Medicine, St. Louis, MO, USA, ^3^ Departments of Laboratory Medicine and Immunobiology, Yale School of Medicine, New Haven, CT, USA, ^4^ School of Biotechnology and Biomolecular Sciences, The University of New South Wales, Sydney 2052, New South Wales, Australia

¶ These authors contributed equally to this work. SF and TEA are Joint Authors

^#^ Present address: Department of Microbiology, Biomedical Discovery Unit, Monash University, Melbourne, Victoria, Australia

Keywords: Mouse Norovirus; Integrated Stress Response; Protein translation; eIF2α; Stress granules; Protein kinase R

Running title: MNV arrests host protein translation

This file includes: Methods, Results and Figure legend for Supplementary Figures 1 and 2

**Materials & Methods**

**Secreted Luciferase reporter assay**

The Ready-To-Glow™ Secreted Luciferase Reporter System assay (Clontech) was performed according to the manufacturer’s protocol. Briefly, cells were transfected with the pBI-CMV5 vector containing the secreted *Metridia* Luciferase sequence alone or an additional sequence encoding one of the MNV proteins or mCherry. 24 h after transfection, 50 µL of the culture supernatants (secreted Luciferase) or the lysates (intracellular Luciferase) were transferred into white bottom 96-well microtest plates (BD Biosciences), before 5 µL of the Luciferase reagent was added. The luminescence was measured directly with the FLUOstar® Omega or CLARIOstar® plate reader (BMG Labtech) and the ratio of the intracellular to extracellular amount of *Metridia* Luciferase activity was calculated.

**Results**

**PABP expression is unaffected by MNV infection.** To explore other regulators of host cell translation, we expressed PABP-GFP in RAW264.7 cells and infected cells for 12h or left them uninfected before cells were fixed and analysed via IFA (Figure 6 A). In mock infected cells PABP-GFP shows a cytoplasmic and rather diffuse staining, which is unchanged in MNV infected cells, which were identified through positive staining with an anti-NS6 antibody. Neither the viral protein NS5 (Suppl. Figure 3) nor NS6 co-localised with PABP-GFP, indicating a lack of interaction between the proteins. Furthermore, we investigated the potential effect of the NS proteins NS3, NS6 and NS7 on the expression of PABP. 293T cells were transfected with PABP-GFP, NS3, NS6 or NS7 and co-transfected with PABP-GFP and the single NS proteins. Expression levels of PABP-GFP and the NS proteins was analysed via immunoblotting (Figure 6 B). PABP-GFP and the NS proteins were detected in high amounts when expressed on their own. Upon co-transfection with NS6 and NS7, PABP-GFP expression levels seemed unperturbed, showing similar intensities for PABP-GFP compared to the control. It is important to note that we were unable to detect a change in size for PABP-GFP or the occurrence of an additional PABP-positive band, indicating that neither of the tested NS proteins seems to cleave PABP. Interestingly, PABP-GFP levels were reduced in PABP-GFP and NS3 expressing cells, possibly due to the detrimental effect of NS3 on translation.

**MNV infection does not impede constitutive protein secretion.** To test if the low amount of secreted cytokines from MNV infected cells is due to a block in protein secretion, we used the bidirectional reporter vector pBI-CMV5, which encodes a secreted *Metridia* luciferase (*Met*Luc). We introduced a fluorescence marker (mCherry) into the vector to be able to identify pBI-CMV5-positive cells. RAW264.7 cells were transfected with the pBI-CMV5-mCherry vector, sorted for mCherry-positive cells via FACS and were either infected with MNV, treated with BFA or left untreated (Figure 17 A). Cell culture supernatants were analysed for their luciferase activity after 12 h of h infection. Transfected cells treated with BFA showed lower activity for *Met*Luc in the supernatant, because the ER to Golgi transport is blocked by BFA. In contrast to that, untreated cells transfected with pBI-CMV5-mCherry secreted high amounts of *Met*Luc into the supernatant. When cells were transfected with pBI-CMV5-mCherry and additionally infected with MNV, there was no significant difference between infected and uninfected cells. Similar levels of *Met*Luc were secreted into the supernatant of infected cells, suggesting that MNV is not influencing the general protein secretion of the host cell.

In addition to this, we tested the effect of the NS proteins on cellular protein secretion (Figure 17 B). For this, we introduced the MNV NS proteins and VP1 into the pBI-CMV5 vector. 293T cells were transfected with these vectors and analysed for *Met*Luc activity in the cell supernatant and *Met*Luc levels within the cell. The ratio between intracellular and extracellular activity of *Met*Luc was calculated for all tested constructs. Cells transfected with pBI-CMV5 were positive for the secretion of *Met*Luc into the supernatant, whereas pBI-CMV5 transfected and BFA treated cells showed a block in secretion and only low amounts of *Met*Luc in the supernatant while luciferase levels within the cells accumulated. None of the non-structural proteins nor VP1 seemed to affect the secretion of *Met*Luc as all tested MNV proteins had a similar supernatant to intracellular luciferase activity ratio compared to pBI-CMV5 transfected cells.

**Figure Legend**

**Figure S1. Host translation protein PABP is not cleaved during infection with MNV. (A)** RAW 264.7 cells were transfected with a cDNA expression plasmid encoding PABP-GFP for 12hrs and then subsequently infected with MNV (m.o.i. of 5) for an additional 12 hrs. Cells were fixed, permeabilised and stained with antibodies against MNV NS5 (red), MNV NS6 (blue) and PABP-GFP is visualised in green. Samples were analysed via confocal microscopy on a Zeiss 710 and images collated in Adobe Photoshop. **(B)** HEK 293T cell were transfected with cDNA expression plasmids encoding the MNV NS3, NS6 or NS7 proteins tagged with 6xHis epitope and PABP-GFP for 18h. Cell lysates were subsequently collected and analysed via immunoblotting and proteins visualised with anti-6xHis and anti-GFP antibodies. GAPDH was visualised with anti-GAPDH an utilised as a loading control.

**Figure S2. MNV does not affect general protein secretion.** (**A)** RAW264.7 macrophages were transfected with the *Metridia* luciferase containing pBI-CMV5-mCherry vector. mCherry positive cells were sorted and infected with MNV, treated with BFA or left untreated. The relative luciferase activity was measured at 12 h.p.i. (n=3, average +/- SEM, ns:p>0.05, *p<0.05, **p<0.01). (**B)** HEK 293T cells were transfected with pBI-CMV5 vectors containing the individual MNV NS proteins. As controls pBI-CMV5 only and pBI-CMV5 and BFA treated cells were used. Supernatants and lysates were collected 24 h post transfection and the ratio between intracellular (lysate) and secreted (supernatant) luciferase activity was calculated (n=3, average +/- SEM, ****p<0.0001).
