## Supplementary Figue 1 for "Mouse Norovirus infection arrests host cell translation uncoupled from the stress granule-PKR-eIF2α axis"

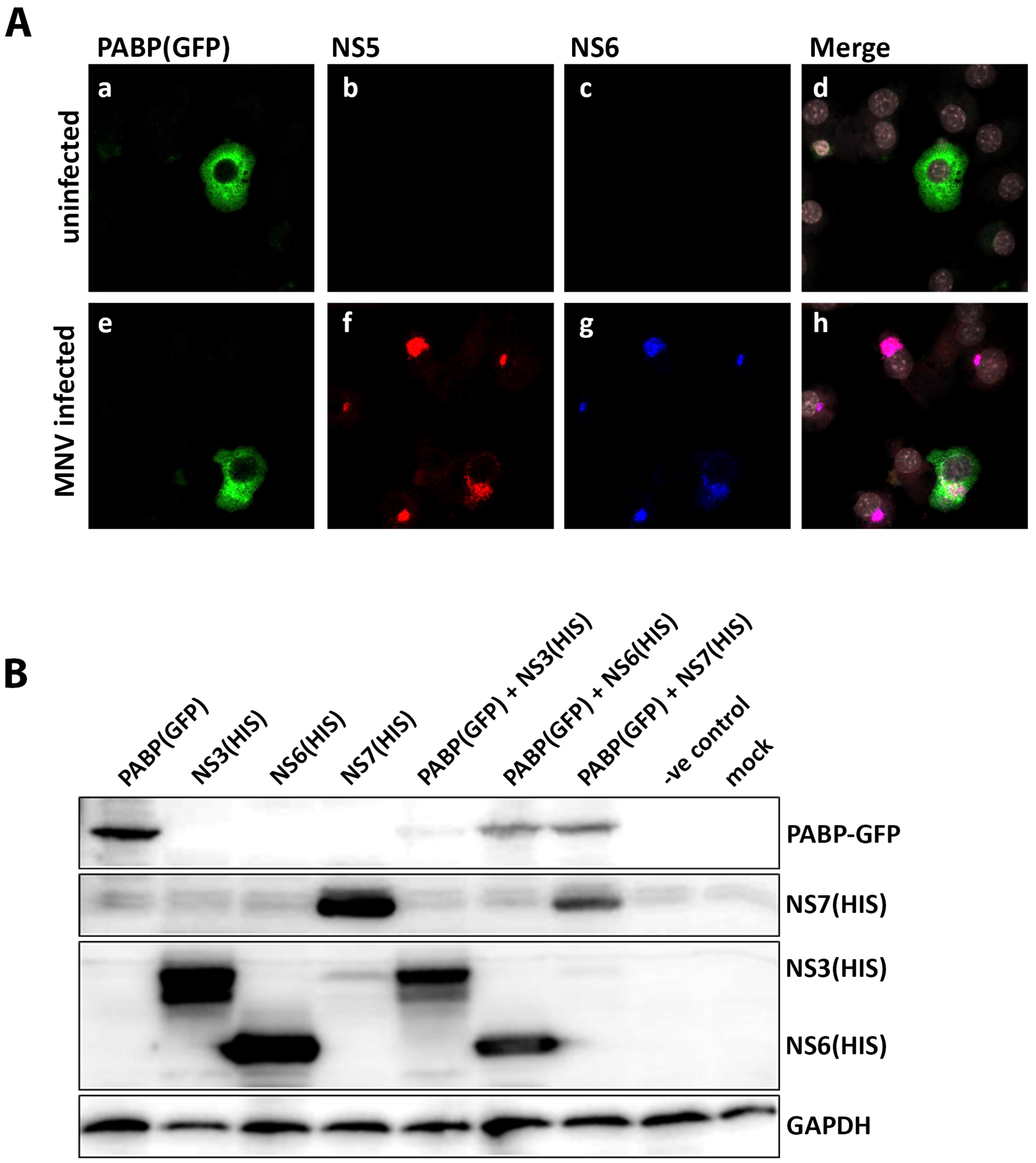


Suppl Figure 1: **(A)** MNV infection does not effect PABP levels however **(B)** NS3 expression reduces PABP levels

Fritzlar, Aktepe et al
