## Supplementary Figue 2 for "Mouse Norovirus infection arrests host cell translation uncoupled from the stress granule-PKR-eIF2α axis"

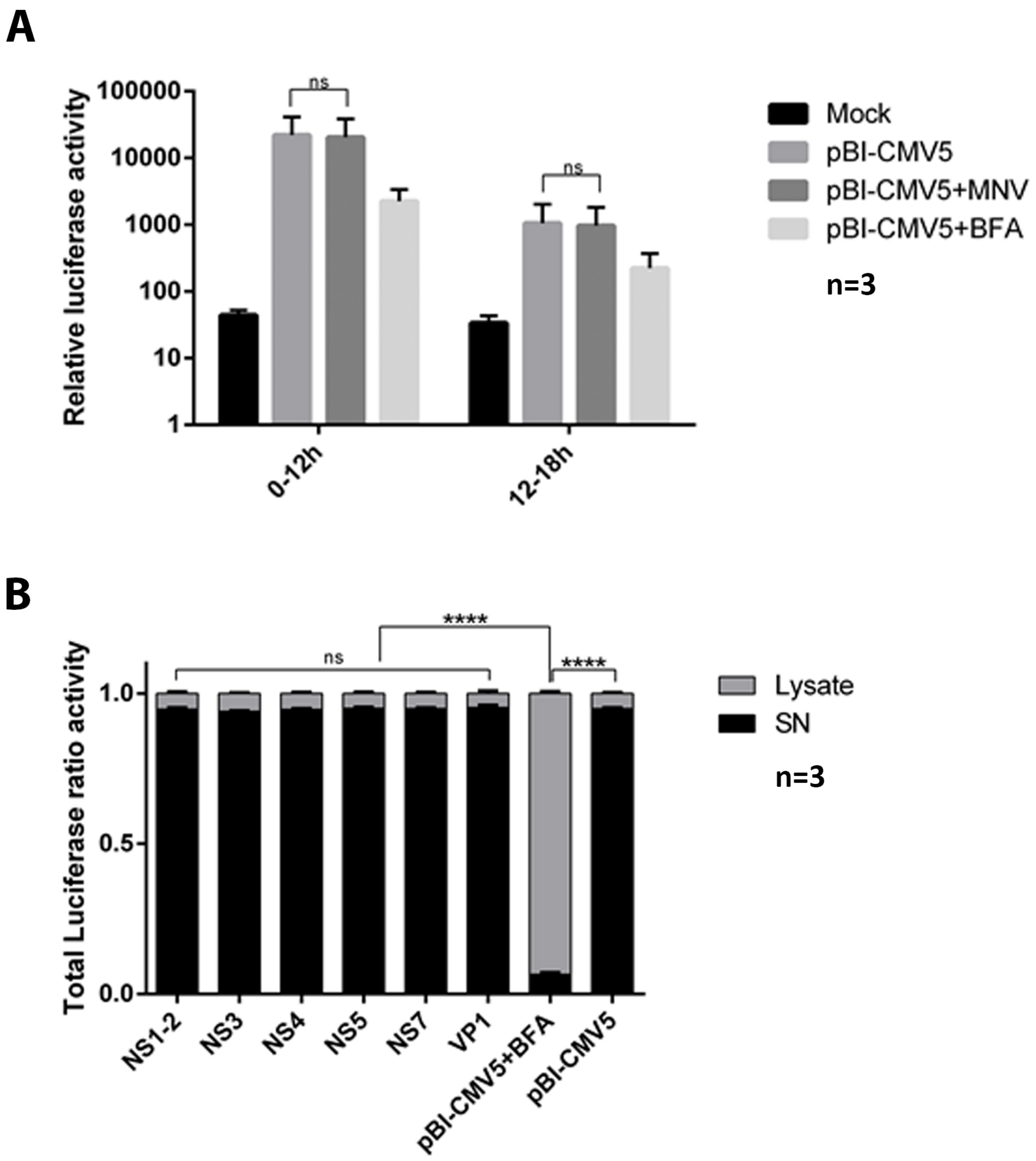


Suppl Figure 2: MNV infection and expression of individual viral proteins does not effect general protein secretion.

Fritzlar, Aktepe et al
